# Hydrogen peroxide modulates lignin and silica deposits in sorghum roots

**DOI:** 10.1101/2021.02.01.429181

**Authors:** Nerya Zexer, Rivka Elbaum

## Abstract

Hydrated silica (SiO_2_·nH_2_O) aggregates in the root endodermis of grasses. Application of soluble silicates (Si) to roots is associated with variations in the balance of reactive oxygen species (ROS), increased tolerance to a broad range of stresses affecting ROS levels, and early lignin deposition. In sorghum (*Sorghum bicolor* L.), silica aggregation is patterned in an active silicification zone (ASZ) by a special type of lignin. Since lignin polymerization is mediated by ROS, we studied the formation of root lignin and silica under varied conditions of ROS by modulating hydrogen peroxide (H_2_O_2_) concentration in the growth solution. Sorghum seedlings were grown hydroponically and supplemented with Si, H_2_O_2_, and KI, a salt that catalyzes H_2_O_2_ decomposition. Lignin and silica deposits in the endodermis were studied by histology, scanning electron and Raman microscopies. Cell wall composition was quantified by thermal gravimetric analysis. We found that the endodermal H_2_O_2_ concentration regulated the extent of ASZ lignin deposition along the root, but not its patterning in spots. Our results show that ASZ lignin is necessary for root silica aggregation in sorghum, and that silicification is enhanced under oxidative stress as a result of increased deposition of the ASZ lignin.

**One sentence summary:** Lignin with carbonyl modifications is patterned by the activity of H_2_O_2_ to nucleate silica aggregations in sorghum roots.

## Introduction

Silicon oxides fertilization is associated with alleviation of biotic and abiotic stresses in an environmentally friendly manner (Liang *et al.*, 2007; Frew *et al.*, 2018; Coskun *et al.*, 2019). Amorphous silica is the most readily dissolved form of silicate minerals (Schaller *et al.*, 2021). Silicic acid molecules in the soil solution are available for plant root uptake through dedicated transporters (Mitani-Ueno and Ma, 2021). Their concerted expression causes super saturation of silicic acid in the xylem sap (Sakurai *et al.*, 2015) of wheat, rice, sorghum and other grasses (Casey *et al.*, 2004; Mitani *et al.*, 2005; Soukup *et al.*, 2020). The silicic acid is transported to the shoot by the transpiration stream and deposited as biogenic opaline silica (SiO_2_·*n*H_2_O). This may lead to accumulation of up to 10% silica per tissue dry weight (Ma and Yamaji, 2006). The deposition takes various mechanisms that are mostly unknown (Hodson, 2016). Heavily silicified locations may be adjacent to cells that are not silicified at all (Lawton, 1980; Bennett, 1982), suggesting a biologically-controlled process. Silica may also form in leaves and roots grown hydroponically without added silicic acid (Kumar *et al.*, 2017; Zexer and Elbaum, 2020), suggesting that biological entities concentrate residual silicic acid to saturation level at the silicification loci.

Recently we reported the first protein that causes silica formation in plants (Kumar *et al.*, 2020). Siliplant1 (Slp1) is expressed in sorghum (*Sorghum bicolor* L.) epidermis cells called silica cells. It is exported to the cell wall, where it interacts with supersaturated silicic acid solution. Slp1 catalyzes the formation of a thick silica wall within hours (Kumar *et al.*, 2017). In parallel, the cell cytoplasm shrinks into threads, and is eventually evacuated during a programmed cell death (Kumar and Elbaum, 2018). Silica is also found in sorghum roots at very specific locations in the endodermis inner tangential cell walls (ITCW), (Sangster and Parry, 1976; Lux *et al.*, 2003; Soukup *et al.*, 2017). However, no Slp1 is expressed in sorghum roots (Kumar *et al.*, 2020). In contrast to leaves, silica in roots is aggregated on densely deposited cell wall, enriched with phenolic material with peculiar characteristics (Zexer and Elbaum, 2020). These phenolic deposits auto-fluoresce in blue (Soukup *et al.*, 2014). Raman microspectroscopy together with chemical manipulations of the tissue indicate the presence of lignin at the aggregation loci (Soukup *et al.*, 2017; Zexer and Elbaum, 2020)

In our previous work, we identified an active silicification zone (ASZ) in roots of sorghum seedlings at about 4-8 cm from the root tip (Figure 1). When plants are grown hydroponically with no Si added, fluorescently green material is deposited at the ASZ, forming spots of about 3 μm in diameter, on a background of non-modified fluorescently blue lignin. With exposure to silicic acid, silica aggregates in the very same fluorescently green locations. The deposition of silica in these spots is associated with a change in their auto-fluorescence from green to blue (Zexer and Elbaum, 2020).

**Figure 1.**
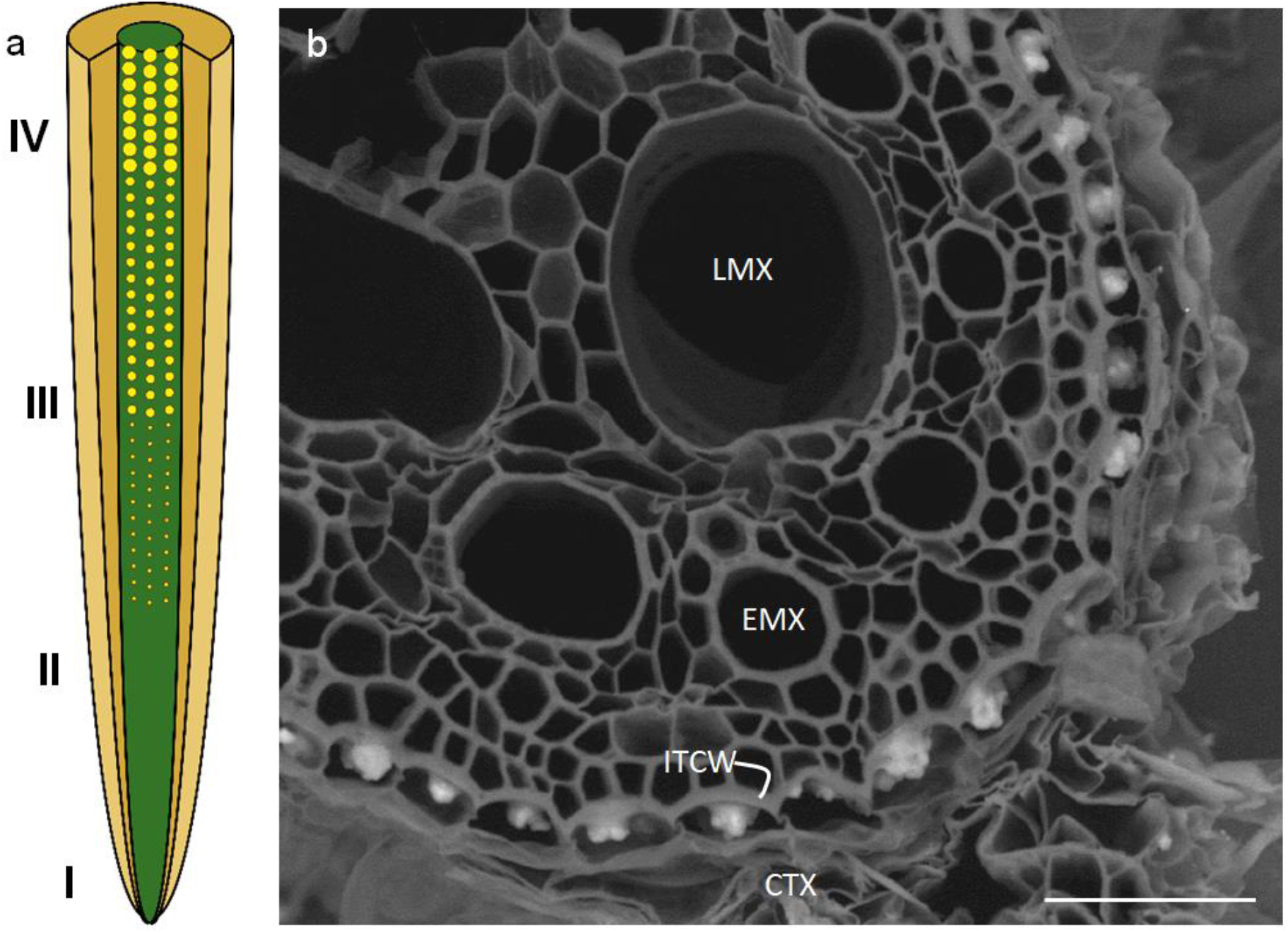
Silica aggregation along the endodermis of a sorghum (Sorghum bicolor L.) seedling primary root. (a) Schematic diagram of a seminal root, drawn without lateral roots (not to scale). In green we represent the central cylinder and in brown the cortex. The endodermis is a single cell layer that resides between these two tissues. Silica aggregates (yellow circles) develop along the root maturation gradient on the inner tangential cell wall (ITCW) of the endodermis. No aggregates are found at the young root tip and up to about 4-6 cm (region I). Aggregation is initiated at about 4-5 cm from the root tip (region II). The aggregates grow in diameter and height along their maturation in regions III and IV. (b) Scanning electron micrograph of a cross section in mature region IV of a sorghum seedling primary root. The central cylinder shows early (EMX) and late meta-xylem (LMX) vessels. The cortex (CTX) was mostly removed. The thick ITCW (ITCW) is indicated. Highly bright silica aggregates are embedded in the ITCW. Scale bar represents 50 μm.

Blue shift in lignin auto-fluorescence indicates a decrease in the system of conjugated pi bonds, possibly as a result of less compact packing of the aromatic rings (Donaldson *et al.*, 2010; Xue *et al.*, 2016, 2020; Donaldson, 2020) or chemical modifications on the rings. The composition and structure of lignin are influenced by the oxidative state at the cell wall during the polymerization of the lignin monomers (Dixon and Barros, 2019; Cesarino, 2019). Specific stressors affect lignification location and extent through the alteration of reactive oxygen species (ROS) (Ali *et al.*, 2006; Denness *et al.*, 2011).

In this work, we tested the hypothesis that the redox balance of the root cell wall influences ASZ lignin formation and the aggregation of silica. For this, we tested silica aggregation in roots of sorghum seedlings under altered redox balance through either stressors that increase H_2_O_2_ apoplastic concentration or KI that quenches endogenous H_2_O_2_. Quantification and mapping of lignin and silica aggregation show that the ASZ lignin, which is needed for silica aggregation, is deposited via the activity of peroxidases and H_2_O_2_, while background lignin is deposited independently of H_2_O_2_ and does not affect silica aggregation.

## Materials and Methods

### Plant material and growth conditions

Caryopses of *Sorghum bicolor* (L.) Moench, line BTx623 and BTx623 *bmr6* mutants were surface sterilized with 3% sodium hypochlorite for 10 min and rinsed twice with distilled water. The *bmr6* genetic stock is PI639713, obtained from the USDA. Grains were then placed in petri dishes lined with wet filter papers and germinated for 72 h. After germination, seedlings were grown hydroponically in non-aerated solutions for 3 days (unless indicated otherwise), after which they were moved to a fresh solution for another 3-4 days. All growth solutions were based on double distilled water containing 1 mM CaCl_2_. Baseline Si concentration was estimated to be 0.01 mM (Nissan *et al.*, 2019). Si+ medium was supplemented with sodium silicate (Na_2_SiO_3_) at a final concentration of 2 mM. Si-medium was supplemented with NaCl at a similar final concentration in order to maintain equal ionic balance across all media. The final pH of all the solutions was adjusted to 5.8 with HCl. In relevant experiments, NaCl (50 mM), KI (1 mM), and H_2_O_2_ (5-50 mM) (Sigma-Aldrich), were added to the solutions.

Seedling were grown on plates of polystyrene foam mounted on 1 L beakers containing growing media and impervious to light. In each beaker we grew ten seedlings. Cultivation was done in a growth chamber, under controlled conditions with photoperiod 16 h : 8 h (light : dark) illuminated with photosynthetic active radiation (PAR) of approximately 200 μmol m^−2^ s^−1^, at a temperature of 28°C : 22°C (light : dark), and 70% air humidity.

### Root tissue fixation and preparation of peeled roots

Complete primary roots were harvested and fixed in ethanol : acetic acid (9:1 v/v) under vacuum for 24 hours. Root samples were kept in the same fixative at 4°C for additional 48 hours. The fixing solution was then replaced with ethanol 70% and samples were stored at 4°C until use. For the preparation of peeled roots, fixed root samples were carefully peeled, exposing the endodermis inner tangential cell wall (ITCW). Removal of cortical tissues was done by finely serrated tweezers (Lux *et al.*, 2003).

### Assessment of the silica aggregate size by scanning electron microscopy (SEM)

Peeled root segments were mounted on aluminum stubs using carbon tape. Observations were performed with a JEOL JSM-IT 100 InTouchScope™ scanning electron microscope (SEM), (JEOL, Japan) under low vacuum (30 Pa) and with accelerating voltage of 20 kV. All micrographs were taken using the back scattered electrons mode. The area in the endodermis inner tangential cell wall (ITCW) occupied by the aggregates was measured by analysis of SEM micrographs using ImageJ2 software (Schneider *et al.*, 2012), as demonstrated in Supplementary Note 1.

### Quantification of the distance from root tip to the active silicification zone (ASZ)

The distance of silica aggregates or lignin spots from the root tip was measured according to the tissue auto-fluorescence (Zexer and Elbaum, 2020). Root length was recorded before peeling. Peeled roots were cut to 1 cm long segments, starting at the root tip and keeping their order up to the root basal end. Segments were then mounted on glass microscope slide using distilled water. The auto-fluorescence pattern of blue spots (in Si+ roots; Soukup *et al.*, 2014) or green spots (in Si-roots; Zexer and Elbaum, 2020) was tracked from the basal section of the roots until no spots could be identified. We calculated the length from the tip of each root to this segment. Observations were performed using a Nikon Eclipse 80i microscope (Nikon, Japan) and an X-cite 120Q (Lumen Dynamics, Canada) UV light source. Specimens were observed using an x20 Nikon lens with the following set of Nikon filters: GFP (excitation: 450–490 nm; emission: 500–550) and DAPI (excitation: 400–418 nm; 450–465 nm).

### 3,3′-diaminobenzidine (DAB) staining and roots sectioning

DAB staining was preformed according to Daudi & O’Brien (2012) with minor changes. Briefly, complete excised fresh primary roots were immersed in petri dishes containing freshly made DAB aqueous solution (5.2 mM, Sigma-Aldrich). Appropriate infiltration of the stain was ensured by applying gentle vacuum for 5 min using a desiccator. Petri dishes were then covered with aluminum foil and placed on a laboratory shaker. After 5 hours of gentle shaking, root samples were transferred to ethanol : acetic acid (9:1 v/v) fixative solution. The solution was replaced once and followed by the fixation procedure described earlier.

Three-millimeter-long root segments were rinsed with double distilled water, positioned in plastic molds, and embedded in optimal cutting temperature (O.C.T) embedding compound (Scigen, USA). The molds were carefully lowered into liquid nitrogen until the O.C.T medium froze. A CM1860 cryostat (Leica) was used to cut 20 μm thick sections, which were mounted on glass slides. These were observed and photographed with a Leica DM500 microscope equipped with a x40 objective coupled to a Leica ICC50W camera.

### Confocal microscopy

Peeled root segments, 10 mm in length, were mounted on glass microscope slides. Samples were immersed in double-distilled water and covered with cover slips. Observations and images were collected using a Leica TCS SP8 confocal laser-scanning microscope equipped with a ×20 objective. The excitation wavelengths used were 405 nm and 488 nm; emission was collected at 400–500 nm and 500–550 nm, respectively. Micrographs were processed using Fiji software and brightness was adjusted evenly across all images.

### Thermogravimetric analysis (TGA)

Peeled primary roots were ground into fine powder using liquid nitrogen and a mortar and pestle (30 roots per sample). The crushed tissue was then dried at 65°C for at least 72 hours before use. Ground and dried samples (16-20 mg) were mounted on platinum pans and measured using a Q500 thermogravimetric analyzer (TA instruments, USA). Pyrolysis was performed in a nitrogen atmosphere and heating ramp of 15°C / min from room temperature up to 900°C. Calculation of the differential thermal gravimetric (DTG) graphs was done using the Universal Analysis 2000 software (TA instruments, USA). The relative quantification of the cell wall polymers was calculated by fitting peaks to the DTG graphs and recording the area underneath the fitted decomposition peaks, using WiRE3.2 software (Renishaw, New Mills, UK).

### Raman micro-spectroscopy

Peeled root segments were mounted on aluminum microscope slides using super glue. The samples were covered with a drop of double-distilled water and measurements were made using an x63 water-immersed objective. Raman maps were collected with a Renishaw InVia spectrometer equipped with a 532 nm laser (45 mW maximum intensity), utilizing WiRE3.2 software (Renishaw, New Mills, UK). Measurements were performed by using the Streamline mode with acquisition time of 30 sec. Maps were composed of between 3000 and 5000 spectra with steps of 1 μm in the x direction. Spectral analysis was done in WiRE3.2 (Renishaw), including peak picking, first derivative calculation, and signal ratio mapping.

### Statistics

Statistical analysis was preformed using the JMP statistical package version Pro15 (JMP, SAS institute Inc., USA). Statistical significance was assumed at *p* ≤ 0.05. For the comparisons of aggregates area, in each treatment at least three roots were used and three micrographs from each root were analyzed (n = 600-1500 aggregates). For measurements of distances form the root tip to the ASZs, at least three roots per treatment were used. The data were subjected to ANOVA followed by the Tukey-Kramer post-Hoc test.

## Results

In order to examine the influence of redox changes in the cell wall on roots silica aggregation, we applied stressors to sorghum seedlings, which are associated with increased endogenous ROS production (Cesarino, 2019). Injury stress was applied to a seminal root of a sorghum seedling grown with no Si supplementation (Si-seedling) by poking with a syringe needle 7 cm from the root tip. The plant was transferred to Si+ hydroponic media for recovery. After 3 days, no aggregates formed at the injury location, possibly due to tissue damage and cell death. In contrast, 0.5 mm above and below the injury, the average area of the silica aggregates was larger in relation to aggregates that developed in parallel regions of an uninjured root (Figure 2, Supplementary Note 1). To corroborate our observations, we grew seedlings similarly under Si-hydroponics, and groups of three plants were moved to Si+ hydroponics and H_2_O_2_ or salinity stress. In these roots, too, we measured larger silica aggregates compared to non-stressed roots (Figure 2, Supplementary Note 1). Interestingly, the increase in the average aggregate area was negatively correlated to the added H_2_O_2_ concentration.

**Figure 2.**
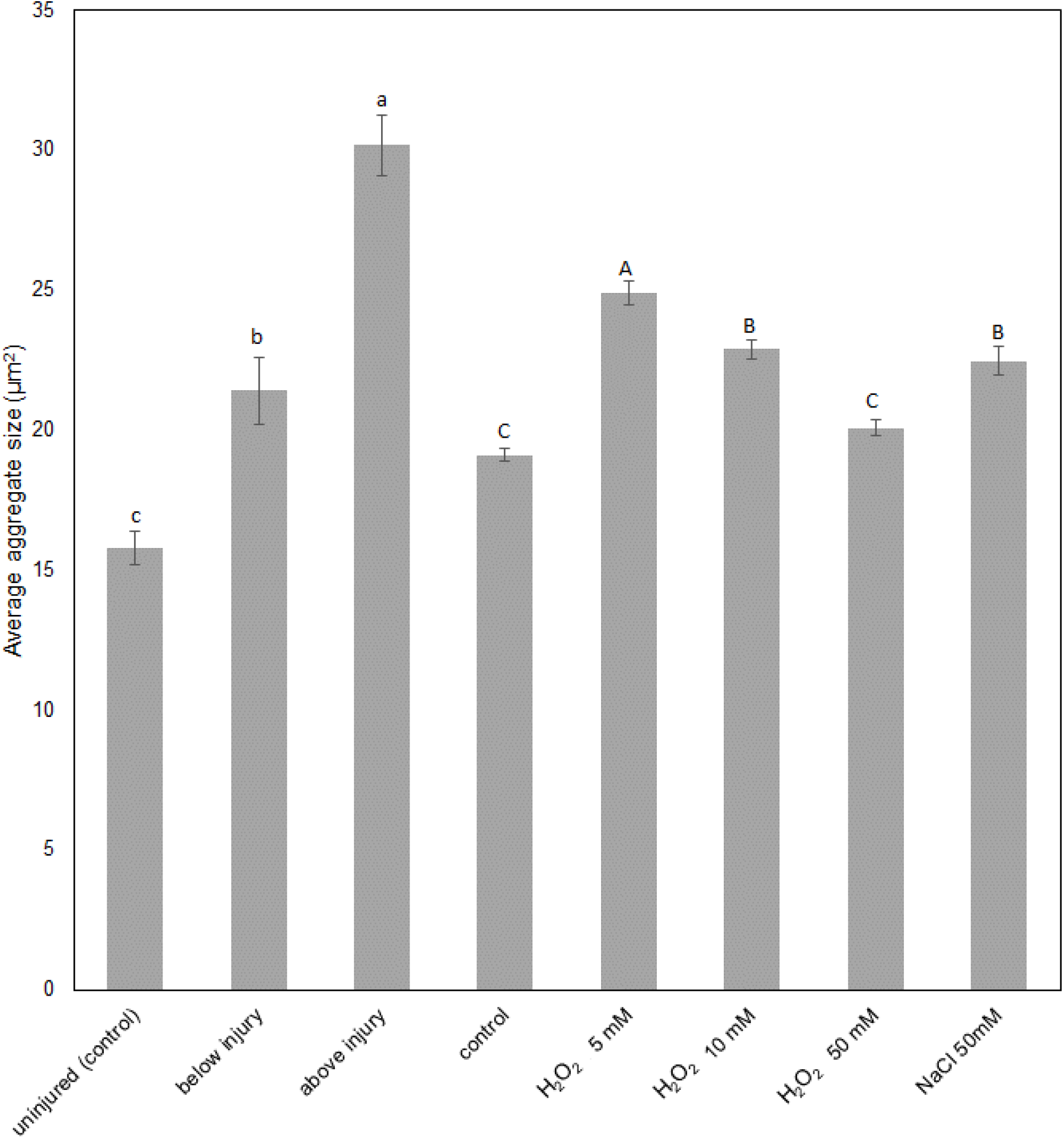
The effect of injury, exogenous H_2_O_2_, and salinity on the area of silica aggregates in the root endodermal ITCW. Presented are means and standard errors. Injury effects were measured 0.5 mm above and below the wound, comparing 500 aggregates from one wounded and one healthy root. Three biological replicates were tested under each of the H_2_O_2_ and NaCl treatments, analyzing at least 500 aggregates in each root. Bars represent standard errors and different letters indicate significant differences at p ≤ 0.05.

To further examine the role of hydrogen peroxide in root silica aggregation, we monitored the activity of peroxidases via 3,3′-diaminobenzidine (DAB) staining. In cross sections of both Si- and Si+ roots, intense staining could be followed from the xylem to the endodermal ITCW. Mapping the peroxidases activity in the ITCW revealed spotted pattern in both Si- and Si+ roots (Figure 3). This indicated a role of peroxidases and H_2_O_2_ in the deposition of ASZ lignin and possibly silica.

**Figure 3.**
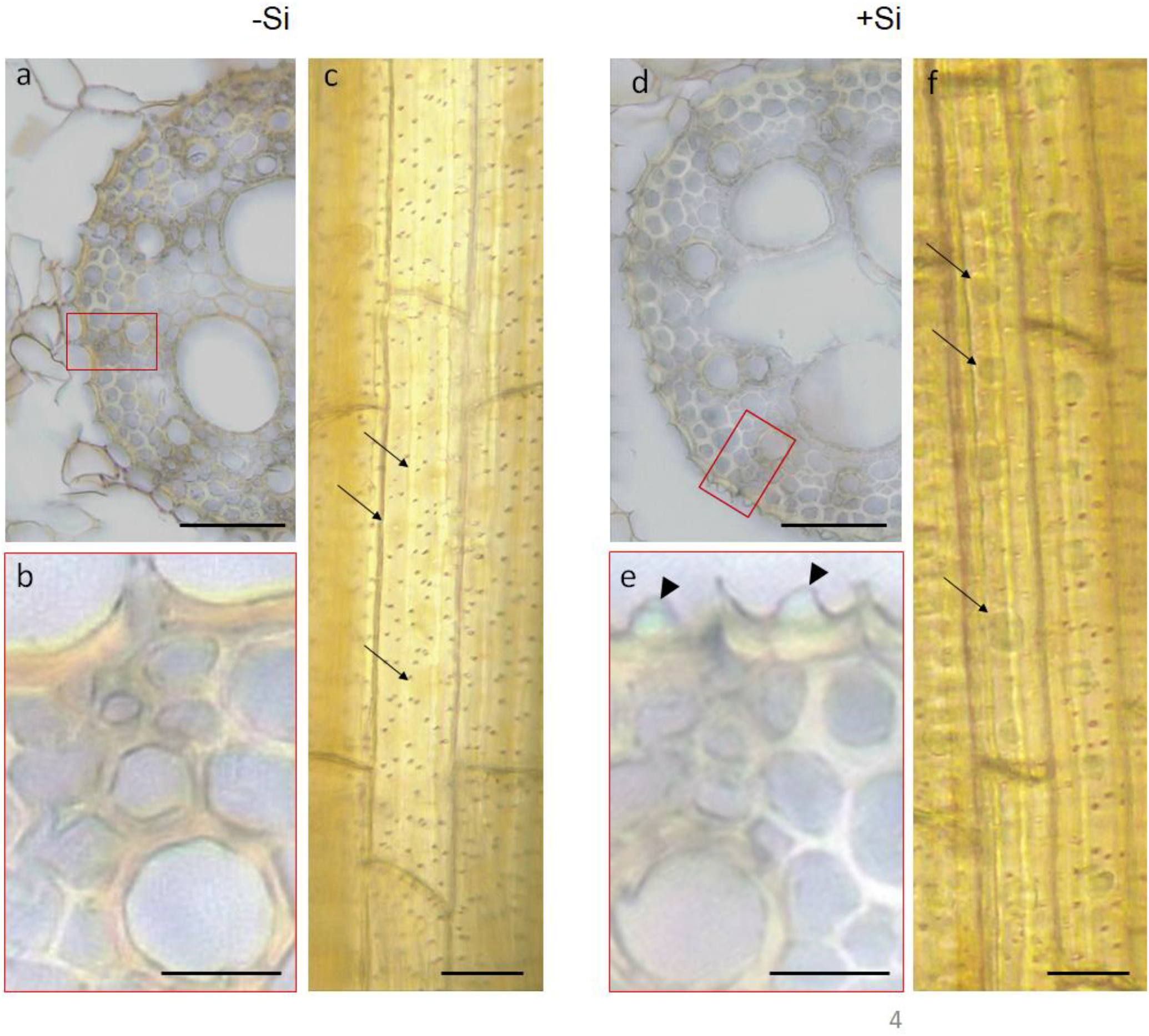
Hydrogen peroxide and peroxidase activity in the primary roots of sorghum seedlings stained with 3,3′-diaminobenzidine (DAB). Cross sections taken from a Si-(a-c) and a Si+ root (d-f). (b) and (e) are magnifications of the rectangles marked in panels a and d, respectively. Arrowheads in e indicate silica aggregates. Brown discoloration attributed to the precipitation of DAB can be seen in the endodermal ITCW and the cell walls of xylem elements. (c) A longitudinal view of DAB stained ITCW from a Si-root. Slightly stronger tainted spots (arrows) can be seen on the background of the stained cell wall. Pits are seen as dark dots. (f) A similar longitudinal view of a Si+ ITCW. The Si aggregates are stained with dark discoloration (arrows). Micrographs are representative of 10 sections taken form five roots of each treatment. Scale bars in a and d represent 50 μm. Scale bars in b,c,e and f represent 10 μm.

To identify the root region in which H_2_O_2_ may affect silica aggregation, we followed silica formation along the root maturation gradient (Figure 1) under high and low H_2_O_2_ levels. For this, seedlings were grown for 3 days in Si-hydroponics solution, allowing a mature non-silicified tissue to develop. On the 4^th^ day, the seedlings were transferred for additional 4 days to Si+ hydroponics solution supplemented with 5 mM H_2_O_2_ or 1 mM KI, which quenched physiological H_2_O_2_. This allowed a younger part of the root to develop under silicic acid and oxidative stress or a reduced native ROS concentration. We then tested these roots for silica aggregation in three regions: (1) young developing tissue about 1 cm from the root tip, developing under altered ROS balance (region I); (2) region composed of cells that were exposed to silicic acid in parallel to completing their maturation under altered ROS balance (region III); and (3) mature region, containing cells that were exposed to silicic acid and altered ROS balance only after completing their development under benign conditions (region IV).

H_2_O_2_ treatment considerably stunted root growth and very likely stressed the roots, while under KI treatment the average length of the roots was comparable to that of untreated samples (Figure 4a-c). In the mature part of the roots (region IV), silica deposited similarly under all treatments (Figure 4d-f). However, silica aggregation was detected already in region I and throughout the root under H_2_O_2_, while under KI, regions I-III were devoid of silica aggregates (Figure 4g-l). When plants were grown similarly in Si-hydroponic solution for 3 days, and then moved for additional 4 days to Si-solution supplemented by H_2_O_2_ or KI, ASZ lignin formed patterns that were similar to the silica aggregation in Si+ roots (Supplementary Note 2).

**Figure 4.**
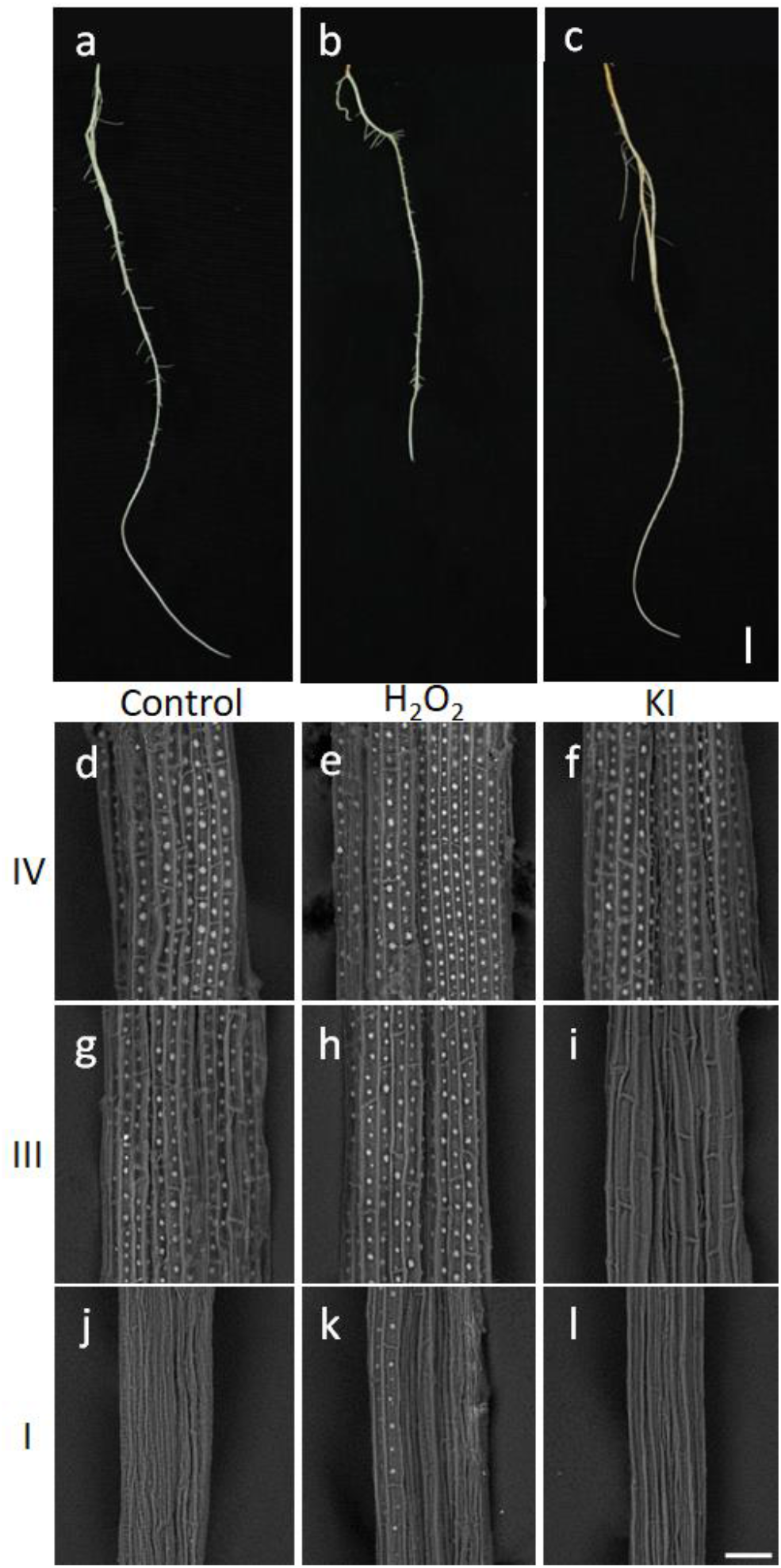
Silica deposition in the endodermal ITCW under high and low H_2_O_2_ regimes. Roots grown in Si-media for three days were transferred to Si+ (control), Si+ & 5 mM H_2_O_2_ (H_2_O_2_), and Si+ & 1 mM KI (KI) treatment media for additional four days. (a-c) Representative images of (a) control, (b) H_2_O_2_, and (c) KI treated roots. (d-l) Scanning electron micrographs of (d,g,j) control roots, and (e,h,k) H_2_O_2_, and (f,I,l) KI treated roots after removal of the cortex. Representative images of region IV, (d-f) region III (g-i), and region I (j-l) are shown. Scale bar in panel c, common to a-c, represents 1 cm. Scale bar in panel l, common to d-l, represents 50 μm.

To quantify the effect of high and low H_2_O_2_ concentrations on silica aggregation (in Si+ roots), in comparison to ASZ lignin formation (in Si-roots), we followed the blue auto-fluorescence of the silica aggregates and green auto-fluorescence of the ASZ lignin (Figure 5 inset). The distance from the root tip to the first auto-fluorescent spot under the various redox conditions was similar between Si+ and Si-treatments (Figure 5).

**Figure 5.**
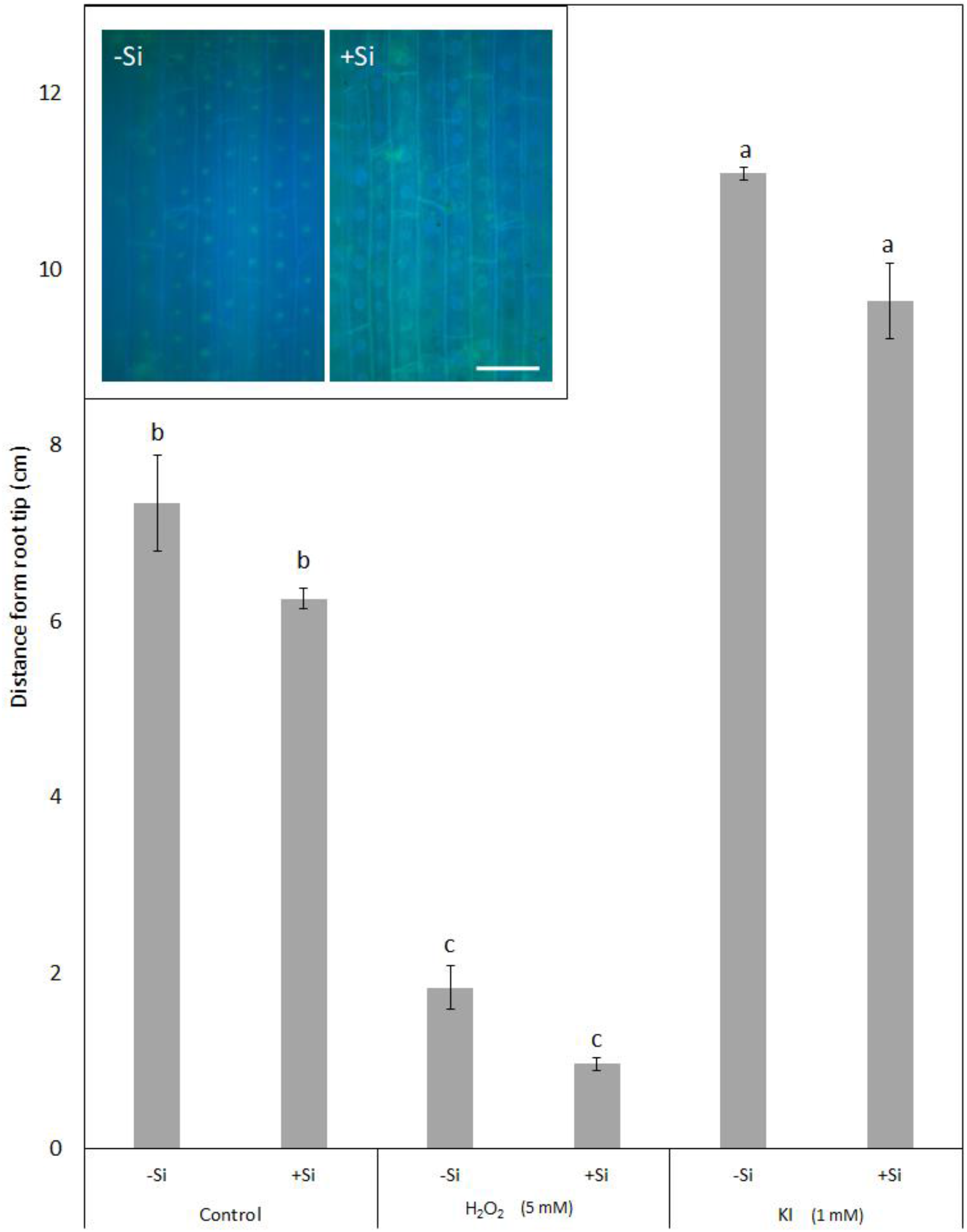
Location along the root of initial silica aggregation (in Si+ roots) or ASZ lignin spotted pattern (in Si-roots). Inset-green and blue auto-fluorescence imaged in the ITCW of roots after the cortex was removed. Images taken from region II of root grown without (Si-, left) and with (Si+, right) Si supplementation under control conditions. Scale bar common to both inset panels represents 20 μm. The average distance from the root tip to the first auto-fluorescent spot is plotted in control, H_2_O_2_, and KI treated roots. In each treatment, five roots were measured. Bars represent standard error. Different letters indicate significant differences at p ≤ 0.05.

Our findings show that the ASZ lignin formation was inhibited by a mild treatment of 1 mM KI solution. We therefore could check whether silica would aggregate in the absence of ASZ lignin. To test this, two treatments were compared. (1) Seedlings were grown under Si- & KI solution for 1 week, and then moved to Si+ & KI for 12 hours. (2) As a control, we grew seedlings in Si-solution, which were transferred after 1 week into Si+ for 12 hours (no KI was applied). In our hydroponics system, the basal 1 cm of the roots closest to the root-shoot junction was surrounded by a polystyrene foam plate that supported the seedlings. Therefore, this region of the roots was not exposed to the growing medium but to air. We monitored silica aggregation in a total of 10 roots in two separate experiments, below and above the hydroponics solution (Figure 6).

**Figure 6.**
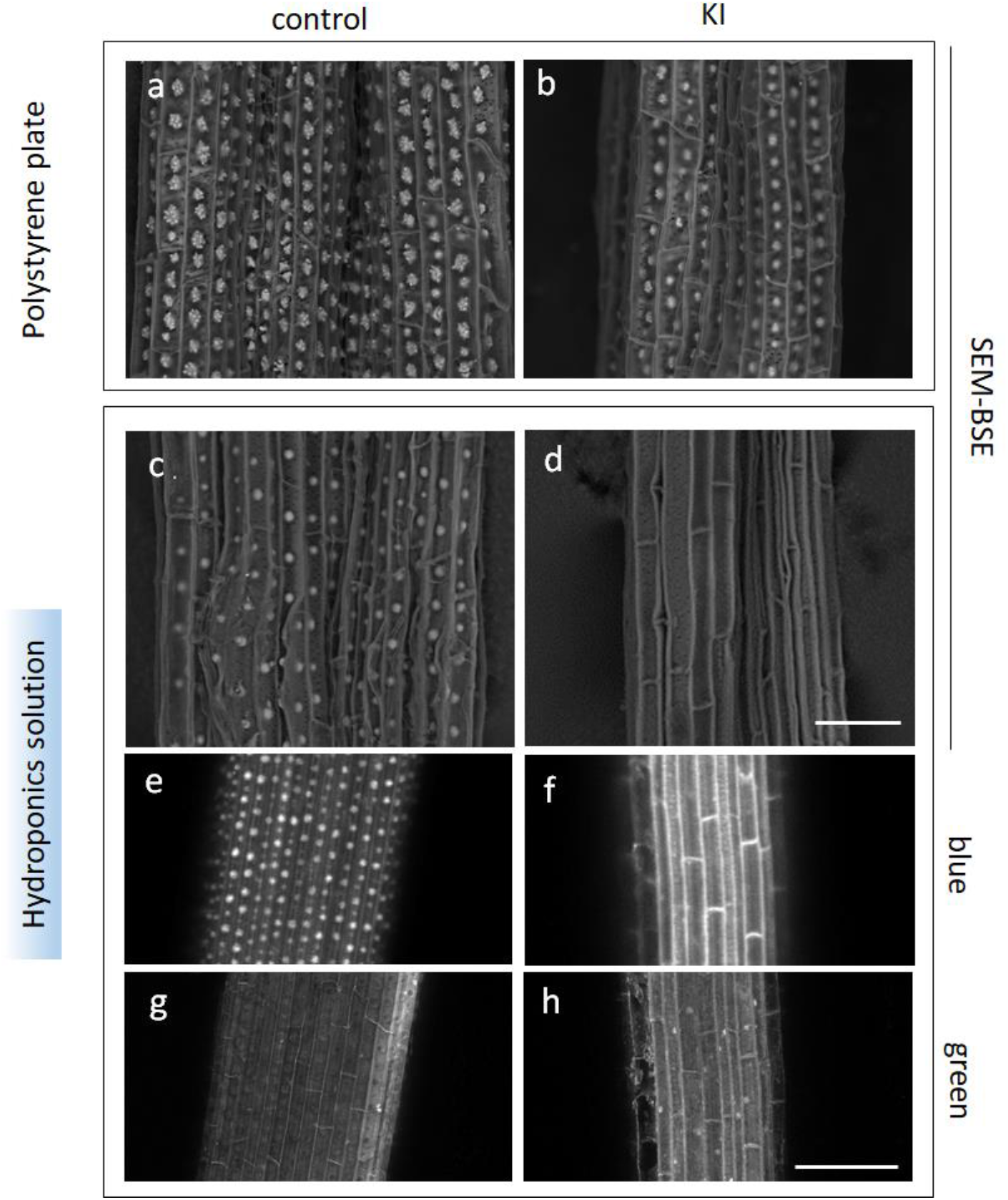
The effect of KI treatment on the formation of silica aggregates. Representative scanning electron micrographs are shown. (a,c,e,g) Roots were grown under Si- for 3 days and then moved to Si+ medium for another 3 days (control). (b,d,f,h) Roots grown in Si-solution were moved to Si+ & KI treatment for another 3 days (KI). (a,b) SEM Micrographs of root sections from the basal end. This root part was not in contact with the hydroponics solutions. (c,d) SEM Micrographs of mature root tissues in region IV that were immersed in the hydroponics solutions. (e,f) Blue auto-fluorescence image of root region IV. (g,h) Green auto-fluorescence image of the same root sections. While blue auto fluorescent patterning of Si aggregates is evident in the control treatment, green pattering is mostly distributed in the background cell wall and around the aggregates. The auto-fluorescent pattering is missing in roots continuously grown in KI. The exposure times for each filter set was similar between the Si treatments. Scale bar in d, common to panels a-d and scale bar in h, common to panels e-h, represent 50 μm.

In the basal root part, never exposed to KI solution, silica aggregates developed in all tested roots, regardless the treatment (Figure 6a,b). In the immersed parts of roots cultivated in KI solution, aggregates were missing altogether (Figure 6d). Spotted pattern of blue auto-fluorescence encircled by green auto-fluorescence developed Si+ roots treated with no KI (Figure 6e,g). This pattern appears normally in Si+ roots (compare to Figure 5 inset, and Zexer and Elbaum (2020)). In roots treated with KI, the blue and green auto-fluorescence was distributed throughout the ITCW and in the radial walls of the endodermis. Fluorescence patterning of spots could not be identified in any of the roots that were continuously treated with KI (Figure f,h). The diffuse blue auto-fluorescence at the endodermal ITCW indicated the presence of some lignin even under quenching of H_2_O_2_.

To estimate the effect of KI treatment on total lignin in the root stele, we quantified the fractions of hemicellulose, cellulose, and lignin by thermal gravimetric analysis (TGA), (Yang *et al.*, 2007; Abidi *et al.*, 2008; Abraham *et al.*, 2018). Three replicates of cortex-peeled primary roots from 30 seedlings were analyzed (Figure 7). Weight losses were measured as a function of the temperature (Figure 7a), and weight loss rate per temperature was calculated (Figure 7b). We deconvolved the rate graph into peaks representing cell wall components, with maxima at maximal decomposition rates, and area representing the fraction per total sample dry weight (Figure 7c). Lignin pyrolysis temperature was similar between the treatments, at 434 ± 44°C. Cellulose decomposed in two stages, at 311 ± 3°C and 323 ± 2°C, similarly in both KI treated and non-treated samples. We detected a reduction in the pyrolysis temperature of hemicellulose in KI treated roots (266.5 ± 0.3°C) as compared to the non-treated roots (283 ± 4°C). Variations in the weight fractions of these three main cell wall components were non-significant at 5% level (Table 1).

**Figure 7.**
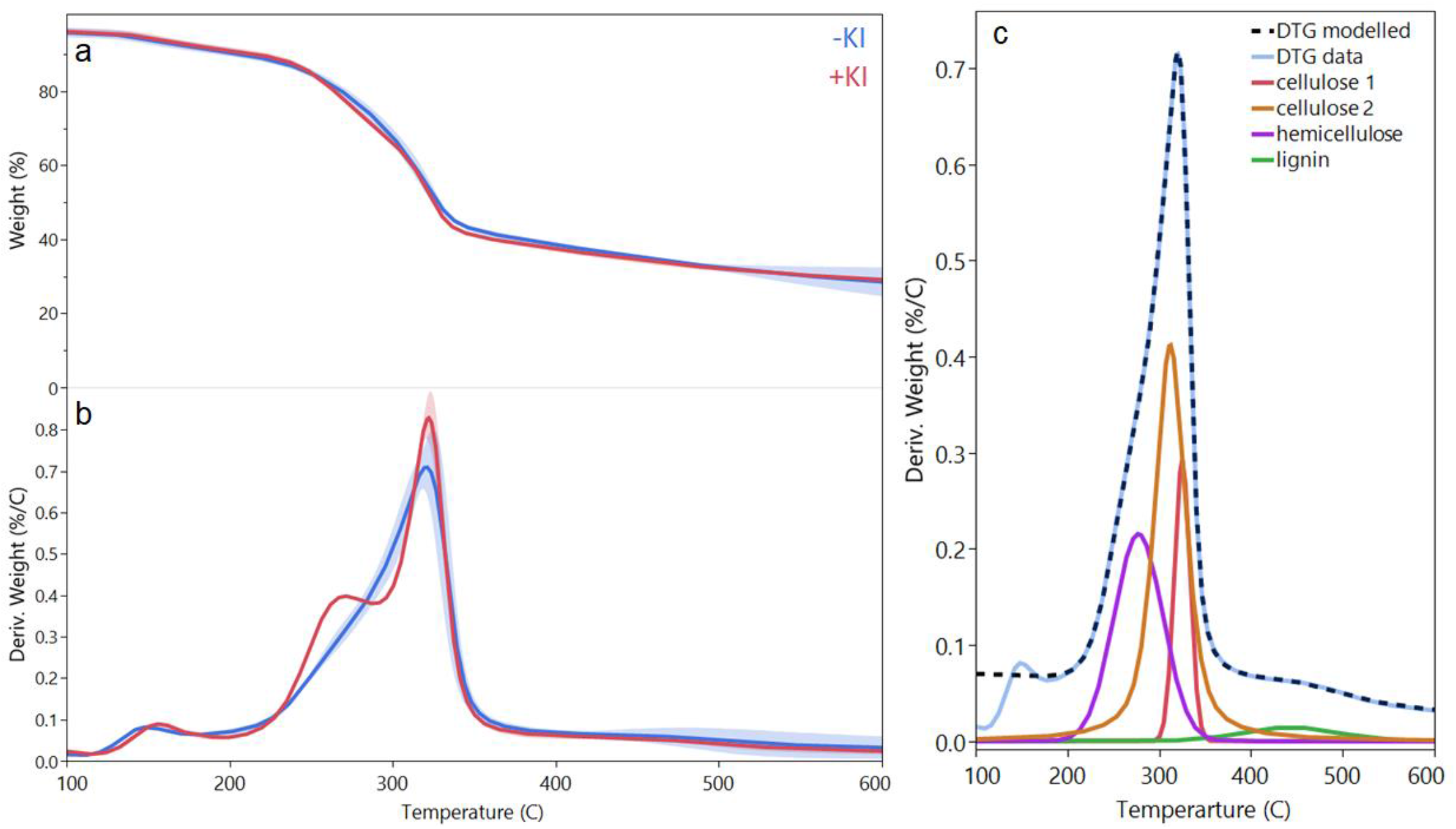
Cell wall composition quantified by thermal gravimetric analysis (TGA) of peeled roots, representing the stele and the endodermal ITCW that are the main locations of lignin deposition in roots. Plants were grown with no Si added, under KI+ (red line) or KI-treatments. (a) Weight loss as a function of temperature of Si-roots grown under control (no KI, blue) or 1 mM KI (red) treatment. (b) Differential thermal gravimetric (DTG) analysis, reporting weight loss rates as a function of the temperature. An average of three biological repetitions of 30 roots with removed cortex are presented for each treatment. The shaded bands represent one standard variation. (c) Example of the calculation of cell wall components through peak deconvolution. A DTG curve (light blue) of one control sample (Si-, no KI) and the modeled DTG curve (black dashed line) calculated by summing fitted curves of cell wall components (red and orange, cellulose; purple, hemicellulose; and green, lignin). Calculated fractions of the cell wall polymers are reported in Table 1.

**Table 1.** Composition of the cell walls of cortex-peeled seedling’s roots based on thermal gravimetric analysis (TGA). Peeled roots were analyzed, because the remaining stele and endodermal ITCW are the main locations of lignin deposition in roots. The average weight percent and standard deviation of hemicellulose, cellulose, and lignin was calculated by peak fitting to the differential weight percent at about 266 – 283°C (hemicellulose), 311 – 323 °C (cellulose), and 390 – 480°C (lignin). Averages of 3 samples each containing 30 roots is reported.

|  | Hemicellulose (%) | Cellulose (%) | Lignin (%) |
| --- | --- | --- | --- |
| **KI-** | 44 ± 11 | 49 ± 13 | 7 ± 2 |
| **KI+** | 32.6 ± 0.8 | 64 ± 1 | 3 ± 2 |
| **P (T-test)** | 0.23 | 0.19 | 0.09 |

We then aimed to identify subcellular variations in the composition of the ITCW in mature root tissues, region IV, as a result of KI treatments. Specifically we aimed to eliminate ASZ lignin by KI treatment, and compare Raman spectra of the endodermis ITCW to a control sample that contains ASZ lignin. All samples were grown with no Si added to avoid variations due to the interaction of the ASZ lignin with silica in the control samples. Figure 8 shows data from a Si- KI- and a Si-KI+ root sample, representing Raman maps collected from about 15 KI-roots, and 3 KI+ roots. Average spectra of circa 3000 spectra in whole Raman maps from both Ki- and KI+ samples showed typical lignocellulosic trace, with major cellulose peaks at 378 and 1095 cm^−1^, and lignin at 1171, 1600, 1630 and 1692 cm^−1^ (Figure 8a), (Agarwal and Ralph, 1997; Agarwal, 2014). Spectral first derivative enhanced hidden peaks and reduced background signals. KI treated samples showed weaker lignin scatterings at 1171 and 1200 cm^−1^ than the control samples. Shoulders at 1585 and 1660 cm^−1^ were identified only in the non-treated control roots (Figure 8b,c). The 1585 cm^−1^ shoulder was assigned to lignin in sugarcane, and the 1660 cm^−1^ shoulder to lignin in wood (Agarwal and Ralph, 1997; Agarwal, 2014). Mapping the total Raman signal before normalization in all non-treated roots revealed a spotted pattern, indicating that the cell wall in these locations is denser than their surroundings (Figure 8d). This was in accordance with our published finding of higher intensity of SEM back scattered electrons at the spots, relative to the surrounding cell wall (Zexer and Elbaum, 2020). In the KI treated samples, no spots were identified, indicating that dense cell wall did not develop (Figure 8e).

**Figure 8.**
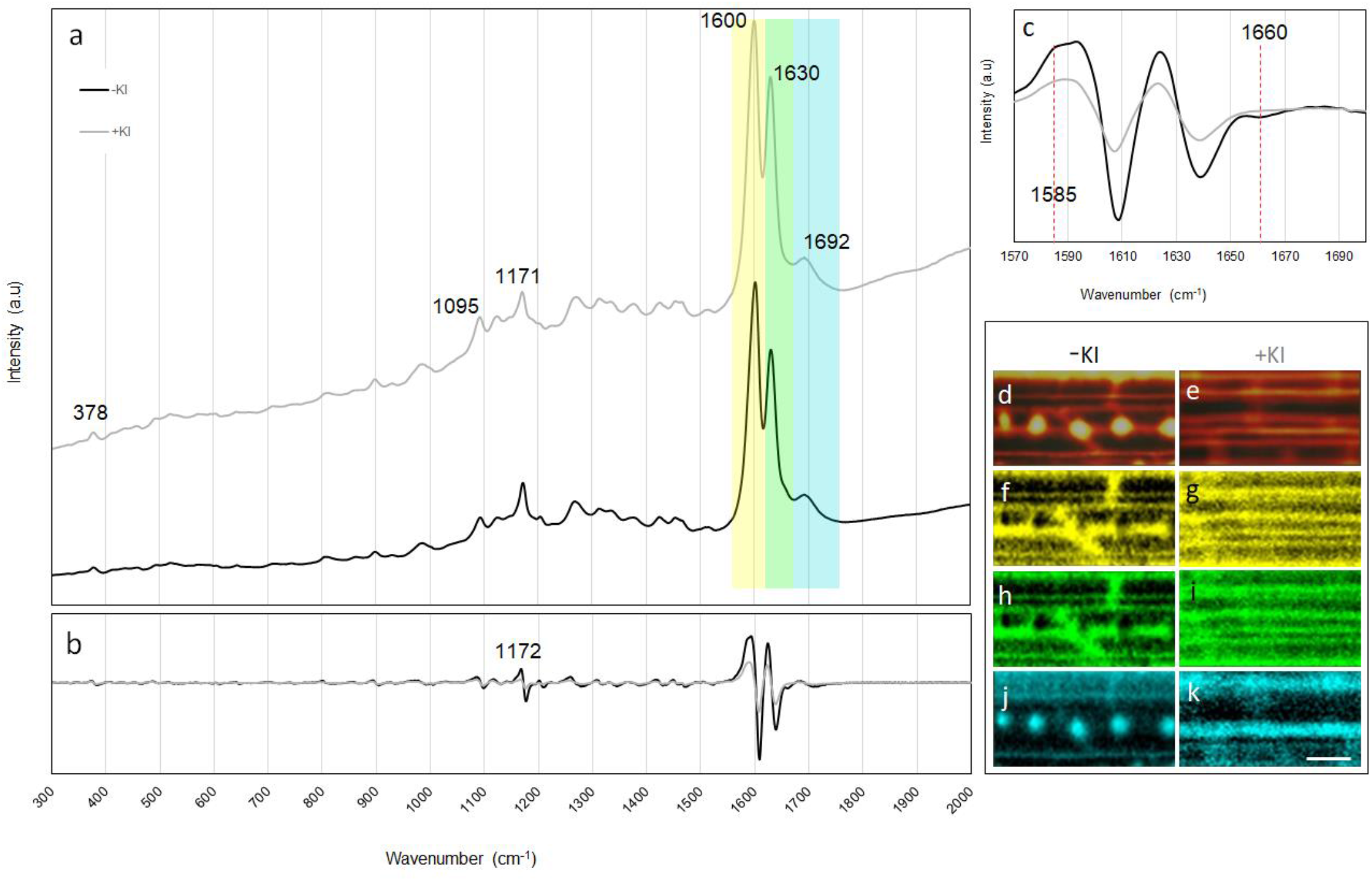
Representative Raman micro-spectroscopy of endodermal ITCW in Si-roots grown under control or KI (1 mM) treatment. (a) Average spectra of ITCW map of control (black, -KI) and KI (gray, +KI) treated roots. The colored rectangles indicate spectral regions mapped in panels f-k, normalized to the crystalline cellulose peak at 378 cm^−1^. (b) First derivative of the average spectra in panel A. (c) Derivative of the spectral range of 1570-1670 cm^−1^ demonstrating the reduction in shoulders at 1585 and 1660 cm^−1^ under KI treatment. (d-e) Maps of the total Raman signal measured from the ITCW of (d) control and (e) KI treated roots. (f-k) Maps depicting typical lignin signals normalized to the crystalline cellulose peak at 378 cm^−1^ of control (left) and KI (right) treated roots. (f-g, yellow) 1550-1618 cm^−1^, (h-i, green) 1618-1660 cm^−1^, (j-k, cyan) 1660-1750 cm^1^. Scale bar in k, common to d-k, represents 20 μm.

In order to identify specific Raman peaks that are related to the spots, while compensating for the general higher signal due to the denser wall, we mapped lignin peaks normalized to the crystalline cellulose peak at 378 cm^−1^ (Figure 8f-k). Interestingly, the aromatic signal at 1660-1750 cm^−1^ was enriched in the spots (Figure 8j), and so was the 1660 cm^−1^ shoulder. This spectral region may indicate the presence of aromatic carbonyls (Agarwal and Ralph, 1997; Taylor and Zografi, 1997). In contrast, the relative intensity of the major lignin doublet at 1590-1610 cm^−1^ and 1620-1630 cm^−1^ did not illuminate the spots (Figure 9f,h), and neither the shoulder at 1585 cm^−1^ nor peaks at 1171 and 1200 cm^− 1^. This indicated that generally, lignin was distributed evenly in the endodermis ITCW, however, the aromatic carbonyl groups were enriched in the spotted pattern of the ASZ lignin. In the roots treated with KI, we could not detect the spotted pattern by any mapping strategy (Figure 8e,g,i,k).

Aromatic carbonyl moieties in cell wall polymers may be found in lignin or suberin. To rule out the presence of suberin at the spots of control roots grown without added Si, we stained the ITCW by Sudan IV. We further compared Raman averaged spectra of Si- KI- roots at the spots to the background cell wall in the spectral region of 2800-3100 cm^−1^. This region shows C-H vibrations that are abundant in the aliphatic part of suberin. Our results showed no evidence for suberin (Supplementary Note 3).

The incorporation of carbonyl residues into lignin is increased in tobacco plants under reduced activity of cinnamyl alcohol dehydrogenase (CAD), (Stewart *et al.*, 1997). Similarly, a mutation in CAD that causes a brown mid-rib phenotype in the *Sorghum bicolor* mutant *bmr6* (Sattler *et al.*, 2009), increases the fraction of carbonyl aldehydes in the lignin (Yahiaoui *et al.*, 1997). To test whether this increased carbonyl fraction may induce increased aggregation of silica, we compared sorghum *bmr6* mutant on the background of line BTx623 to wild type BTx623 sorghum. Three plants of wild type and mutant genotypes were grown hydroponically in Si+ solution for one week. Roots were extracted and peeled for the assessment of average silica aggregate area in region IV (Supplementary Note 1). We measured mean aggregate area (± SE) of 27.8 ± 0.5 μm^2^ in the *bmr6* mutant plants, significantly larger than the average area of 19.4 ± 0.3 μm^2^, measured in wild-type plants (p<0.001).

## Discussion

### Silica deposits require an optimal level of H_2_O_2_

In sorghum roots, silica aggregates are deposited in a neatly ordered pattern as part of the endodermal ITCW. This process is only attained by live tissue and is metabolically controlled (Soukup *et al.*, 2020). Damaging the root locally prevented the formation of aggregates at the site of the wound, possibly as a result of cell death at the endodermis. In healthy tissue near the wound, aggregates were bigger than in non-wounded tissue, and more so above the wound than below it. Our results showed that excess H_2_O_2_ concentration is negatively correlated to the aggregate area, with highest effect at the lowest concentration tested (Figure 2). The complex relationship between ROS balance and silica deposition is in accord with the dual role of ROS in redox biology, that at low concentration induces cell proliferation, differentiation and elongation, and at high concentration signals for stress and cell death, and may apply stress via oxidation of useful molecules (Tsukagoshi *et al.*, 2010; Schieber and Chandel, 2014). Nonetheless, under very low endogenic H_2_O_2_ concentration, during KI treatments, aggregates did not form even in plants supplied with silicic acid (Figure 6). Since the tissue still contained lignin (Figure 6–8), it appears that the general deposition of lignin occurred also under a minimal level of H_2_O_2_. However, the generally deposited lignin was not enough for silica aggregation. We conclude that a minimal level of H_2_O_2_ is required for the formation of ASZ lignin that patterns silica aggregation.

### Low H_2_O_2_ concentration reduces hemicellulose cross-linkage but not lignin

Under KI treatments, the hemicellulose pyrolysis temperature was lower by 16°C (266.5 ± 0.3°C in treated roots and 283 ± 4°C in non-treated roots) (Figure 6). Such a decrease in thermal stability may be a result of reduced levels of crosslinking (Farhat *et al.*, 2017). Further, the increase in variation of hemicellulose pyrolysis temperature, which is 10 times higher in the control samples as compared to the samples grown under KI may result from the random nature of these crosslinks. In contrast, the pyrolysis temperature of lignin was similar between the treatments, suggesting that lignin cross-linkage was not altered (Stewart *et al.*, 1997).

Hemicellulose polymers are crosslinked to one another through di-ferulic acid ester bonds and to lignin by ester or ether bonds of ferulic or coumaric acid (Fry *et al.*, 2000; Hatfield *et al.*, 2017). Reduced incorporation of ferulic acid in the cell wall may be identified as a KI-induced reduction in the Raman peaks at 1172 cm^−1^ assigned to ferulic acid (Soukup *et al.*, 2017), and 1585 cm^−1^ assigned to di-ferulates (Abraham *et al.*, 2018). Since these peaks are not mapped uniquely to the ASZ lignin spots, hemicellulose-ferulic acid complexes are suggested not to be involved in the nucleation of silica (Figure 8). Ferulic acid and arabinoxylan were associated with silica aggregates that were isolated by sulfuric acid digestion of sorghum roots (Soukup *et al.*, 2017). Taken together, we propose that hemicellulose-ferulic acid complexes may associate with silica aggregation but are not part of the nucleation complex at the ASZ lignin spots.

### Silica and ASZ lignin patterning

The ASZ lignin in plants grown without silicic acid appeared in a spotted pattern and reacted to redox changes in a similar manner to the silica aggregation pattern in plants fed with silicic acid (compare Figure 4 and Supplementary Note 2). We showed that the ASZ lignin and silica aggregation co-localize by overlapping the fluorescently green lignin spots with localities that nucleate silica (Zexer and Elbaum, 2020). Here we show that the ASZ lignin deposition was independent of silicic acid. In contrast, in roots treated by KI, no ASZ lignin nor silica deposited (Figure 6). A hierarchical relationship appears, in which H_2_O_2_ is required for the formation of lignin that is needed for silica aggregation.

Lignin formation requires a source of radicals. These could be oxygen molecules processed enzymatically by laccase. Alternatively, NADPH oxidase (called respiratory burst oxidase homologs (RBOH) in plants) produces O_2_^-·^ that are converted to H_2_O_2_ molecules, processed by peroxidases. A radical is transferred by the enzymes to a phenylpropanoid, which is coupled to a growing lignin polymer. In Arabidopsis, lignin deposition is patterned by the formation of enzyme complexes bound to the cell wall under specific developmental contexts, such as lignification of root Casperian strips or xylem elements (Lee *et al.*, 2013; Hoffmann *et al.*, 2020). The enzyme complexes include anchoring motifs, RBOHs, superoxide dismutase (SOD), and laccase or peroxidases. In our system, there was a profound influence of apoplastic H_2_O_2_ concentration on the intensity of localized lignification leading to silicification. This suggests the involvement of peroxidases in the formation of the ASZ lignin spots in the tertiary cell wall.

Nonetheless, the spatial pattern of lignification was not altered with addition of H_2_O_2_, suggesting that the endogenous production of H_2_O_2_ does not create the lignin patterning. Therefore, the pattern could originate from a pattern of peroxidases bound to the cell wall, or by modified monolignols (typical to ASZ lignin) that are in some way bound specifically in spots. Lignification occurs over the whole surface of the tertiary cell wall, with lower deposition in the spots, as seen by mapping the Raman total lignin scattering (Figure 8f,h). This suggests the activity of peroxidases and/or laccases throughout the ITCW. Since total lignin formation was only slightly reduced by KI treatment (Table 1), we may assume that the lignin forming in the seedling root is mostly produced as a result of laccase activity, unaffected by low levels of H_2_O_2_. We may suggest that the ASZ lignin pattern forms by patterning of peroxidases on a background of laccases. However, both laccases and peroxidases would interact with whatever lignin monomers they may find (Tobimatsu and Schuetz, 2019), possibly including the ASZ lignin monomers. Therefore, our results suggest that the spotted pattern of modified lignin either originated from patterned ASZ-phenylpropanoid monomers or required the involvement of dirigent proteins that selectively interact with specific monolignols (Dixon and Barros, 2019; Tobimatsu and Schuetz, 2019). Another option would be the oxidation of the polymerized lignin to produce carbonyl modifications *in situ* at the spot location. However, no experimental evidence supports such a modification mechanism (Dixon and Barros, 2019).

Silicon in plants is studied as a stress-ameliorating agent. Reports encompass species from diverse plant families confronting a range of both biotic and abiotic stresses (Liang *et al.*, 2007; Van Bockhaven *et al.*, 2013; Coskun *et al.*, 2019). Many of these works point to Si-generated effects on plants’ oxidative state including gene regulation, protein abundance and ROS levels (reviewed by Cooke and Leishman, 2016). A specific example in root endodermis shows that the supplementation of silicic acid to the growing medium enhances lignification (Fleck *et al.*, 2011; Lukačová *et al.*, 2013) but not suberization (Kreszies *et al.*, 2020) through modulation of ROS and ROS-dependent enzymes (Fleck *et al.*, 2011). In this work, we show a reciprocal effect: the apoplastic oxidative levels affect the extent of endodermal silicification in roots of sorghum seedlings through mediation of lignin formation. This new direction requires further work to show whether the effects of silicon on plants are induced by its interactions with ROS.

## Author Contribution

Both authors conceived and designed the experiments. N.Z conducted the experiments, and both authors analyzed the data and wrote the manuscript.

The research was funded by the Israeli Ministry of Agriculture & Rural Development, Chief Scientist Office grant 12-01-0016. N.Z is thankful to the Robert H. Smith Foundation for a scholarship.

## Acknowledgements

N.Z is thankful to the Robert H. Smith Foundation for a scholarship. The research was funded by the Israeli Ministry of Agriculture & Rural Development, Chief Scientist Office grant 12-01-0016.

