## Supplementary Notes for "Hydrogen peroxide modulates lignin and silica deposits in sorghum roots"

### Supplementary Note 1: Quantification of silica aggregation through the estimation of their area projected onto the endodermis internal tangential cell wall (ITCW)

Assessment of the silica aggregation was based on the area each aggregate occupied in the endodermis internal tangential cell wall (ITCW), as imaged by scanning electron microscope (SEM) at the back scattered electron (BSE) mode. Roots grown in Si+ solution were peeled to expose the ITCW of their endodermis. Panel (a) shows SEM-BSE imaging. Since Si atoms are bigger than carbon atoms, they scatter more electrons during imaging. As a result, the silica aggregates seem as bright areas on a dark carbohydrate background. In order to quantify the area that each aggregate occupies, SEM-BSE images were inverted and made binary through the application of a threshold, as demonstrated in panel (b). After this manipulation, the aggregates were seen as dark round objects. Panel (c) shows the particles identified by applying ‘Measurement’ function in ImageJ2 software (Schneider *et al.*, 2012). The particle area was limited between 5 and 50 $\mu m^2$, with a minimal circularity index of 0.3. Individual particle area was recorded and averages were compared between treatments. Scale bars in all panels are 50 µm. For the comparisons of aggregate area, in each treatment at least three roots were used and three micrographs from each root were analyzed (n = 600-1500 aggregates).

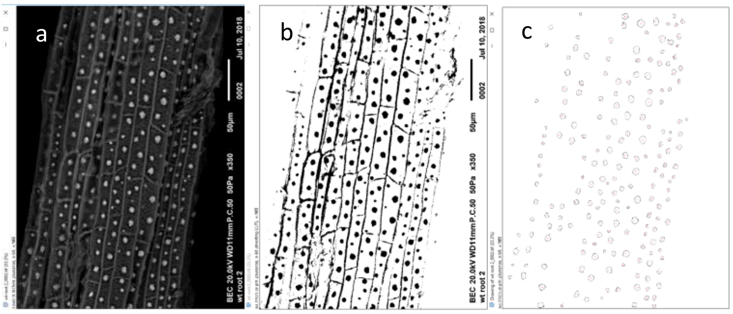

**Supplementary Note 2: Effects of variation in $H_2O_2$ concentration on the formation of ASZ lignin**

Plants were grown in Si- hydroponic solution for 3 days, and then moved for additional 4 days to a solution supplemented by $H_2O_2$ or KI. To visualize lignin and test the formation of ASZ lignin patterns we exposed the endodermis ITCW by peeling off the cortex in 20 fixed roots, and stained root sections with phloroglucinol-HCl. This stain interacts with the 4-O-linked hydroxycinnamyl aldehyde end groups of lignin. Therefore it is considered to be lignin-specific (Pomar et al., 2002). Staining solution was prepared by mixing a 3% phloroglucinol (Sigma-Aldrich) solution in ethanol (100%) (w/v) followed by the addition of 0.5 volume of hydrochloric acid (32%). Desired segments were hand-cut from each root and mounted on glass slides using freshly made staining solution. Slides were immediately observed and photographed with a Leica DM500 microscope equipped with an x40 objective and a Leica ICC50W camera (Leica, Germany).

Panels (a-c) show roots grown in Si- media for three days that were transferred to Si+ media for additional 4 days, and then stained with phloroglucinol-HCl. Unstained silica aggregates are evident on a stained background in regions III (b) and IV (a). There is no staining in region I (c). Panels (d-l) show roots grown in Si- media for three days that were transferred to Si- control, Si- 5 mM $H_2O_2$ and Si- 1 mM KI treatment media for additional four days and then stained with phloroglucinol-HCl. Panels (d), (g) and (j) show micrographs of region IV taken form control, $H_2O_2$ and KI treated roots, respectively. Panels (e), (h) and (k) show micrographs of region III taken form control, $H_2O_2$ and KI treated roots, respectively. Panels (f), (i) and (l) show micrographs of region I taken form control, $H_2O_2$ and KI treated roots, respectively. Arrows point to ASZ lignin spots. Scale bar common to all panels represents 50 µm.

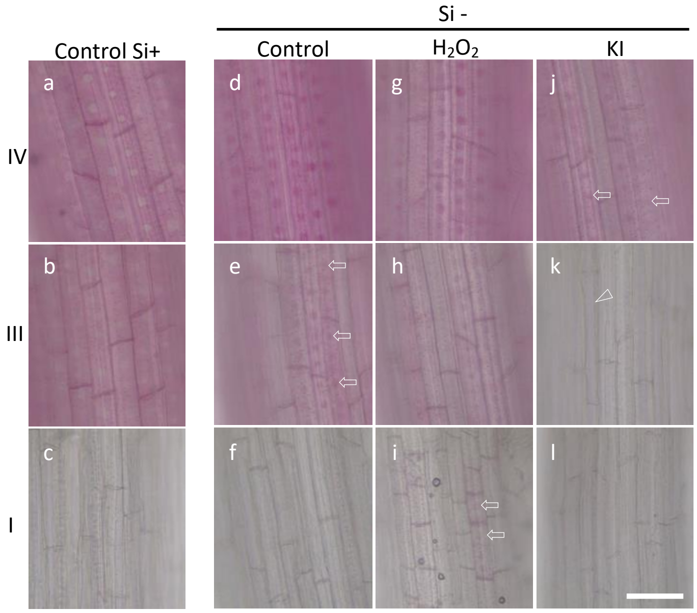


In roots treated with $H_2O_2$, phloroglucinol staining was detected already in region I in spots as well as background coloring (i). A parallel region I did not stain in control Si+ nor Si- roots (c,f). In KI treated roots, staining was radically reduced. No ASZ lignin spots could be found in regions I & III that had completed their development under treatment conditions (k,l). Nonetheless, sporadic faint background coloring in Region III (arrowhead) indicated background lignin deposition (k).

**Supplementary Note 3: Suberin could not be detected at the ITCW spots**

The endodermis is rich in suberin, which may contain aromatic carbonyl moieties, similarly to lignin. In order to test whether suberin is involved in the ASZ lignin structure, we stained root sections by Sudan IV, a non-polar fat-soluble dye. Suberin has lignin-like phenolic domain modified by aliphatic residues. Sudan IV is targeted to the hydrophobic aliphatic domain, where it accumulates and stains suberin red (Lulai and Morgan, 1992).

Sudan IV (Sigma) staining solution was prepared by dissolving 0.25 gr of Sudan IV in 50 mL 95% ethanol. After adding 50 mL glycerol, the solution was mixed and allowed to incubate at room temperature for 1 hour. Roots of sorghum seedlings grown with no added Si were fixed as described in the main text. Hand-cut cross sections and 5 mm long peeled root sections were mounted on microscope slides with a drop of staining solution and covered with a cover slip. After 15 min, samples were observed and photographed with a Leica DM500 microscope equipped with a x20 or x40 objective and coupled to a Leica ICC50W camera.

Panel (a) shows a cross section of the root, and panel (b) a magnification of the marked square in panel (a). Staining throughout the stele was revealed. Intense accumulation of the dye was following a path between the xylem and the endodermal ITCW, reminding the staining of $H_2O_2$ with 3,3'-diaminobenzidine (DAB), see Figure 3 in main text. At larger magnifications (b), suberin lamellae was identified as part of the endodermal ITCW (filled arrowheads). The thin outer tangential cell walls of the endodermis were also darkly stained (voided arrowheads). Staining of peeled roots viewed longitudinally enhanced the residues of radial walls (c). However, we could not identify staining of putative aggregation sites in a spotted pattern. Ctx, cortex; en, endodermis. Scale bars in (a) and (c) represent 50 µm, and in (b) 10 µm.

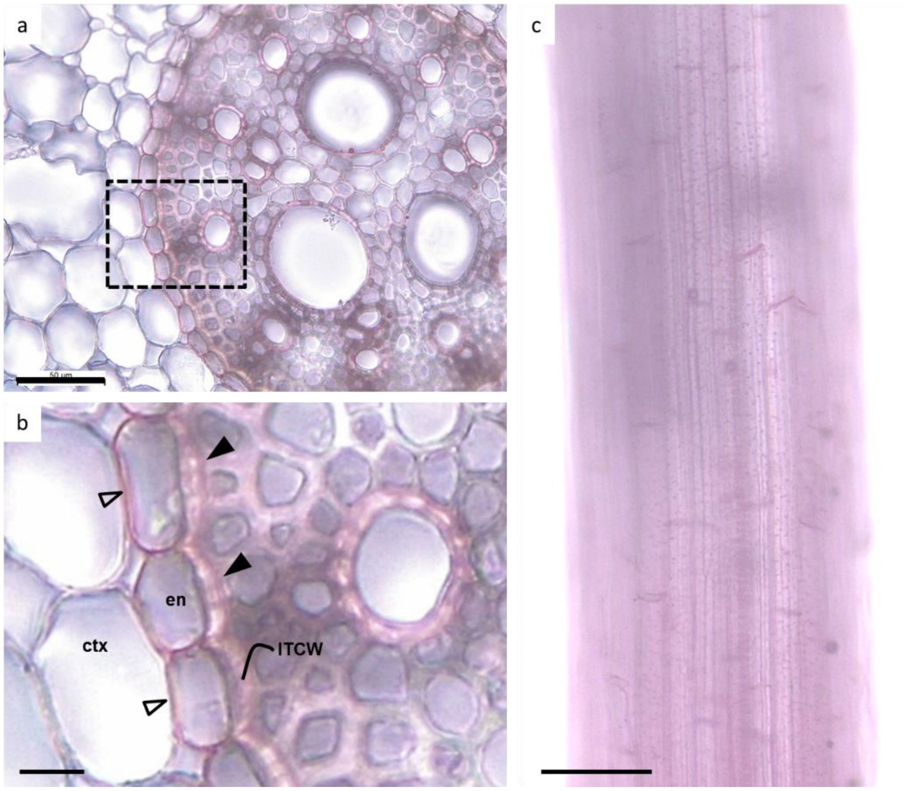

Suberin contains waxy residues (Lulai and Morgan, 1992) that are rich in $CH_2$ chains groups. These have a distinct Raman peak at 2850 and 2880 $cm^{-1}$ (Vulavala *et al.*, 2015; Man *et al.*, 2018). In order to test whether suberin is more abundant in the ASZ lignin spots than in the background ITCW, Raman spectra were extracted from Raman maps. Maps of the ITCW were collected in peeled roots viewed longitudinally, as described in the main text. We averaged at least 20 single spectra from ASZ lignin spots (panels d,e, red lines) and 20 spectra from the background cell wall (panels d,e, blue line). One standard deviation is shown as a faded band following each mean spectrum. Lignin and polysaccharide peaks are indicated in the fingerprint region at wavenumbers below 2000 $cm^{-1}$ (d). A marked difference is the increased background signal at the spots, which is growing considerably above 1700 $cm^{-1}$. This increased baseline is possibly a result of photoluminescence, excited by the 532 mn laser, with a maximum at 2200 $cm^{-1}$, circa 600 nm. In this spectral region we expect overlapping bands assigned to phenols, expected in both suberin and lignin (Vulavala *et al.*, 2015).

To capture Raman scatterings typical to suberin but not lignin we compared the spectral region of 2600-3200 $cm^{-1}$, which includes the C-H vibrations at 2700-3100 $cm^{-1}$. The peaks at 2890 $cm^{-1}$ and 2940 $cm^{-1}$ are assigned to asymmetrical CH stretching vibration of $CH_2$ and the symmetrical CH stretching vibration of $CH_3$, respectively (Anthony, 1982), and were identified in lignin, cellulose, and pectin (Agarwal, 2014). The shoulder at 3070 $cm^{-1}$ was assigned to in plane CH symmetrical stretch of aromatic rings (Larsen and Barsberg, 2010). A shoulder could be detected at 2845 $cm^{-1}$, assigned to lignin (Agarwal, 2014). However, major signals at 2850 and 2880 $cm^{-1}$, typical to suberin, were not apparent in any of our measurements. Attempting to visualize the ASZ lignin spots by mapping the suberin peaks was not possible. Our results reject the hypothesis that suberin is involved in the ASZ lignin structure.

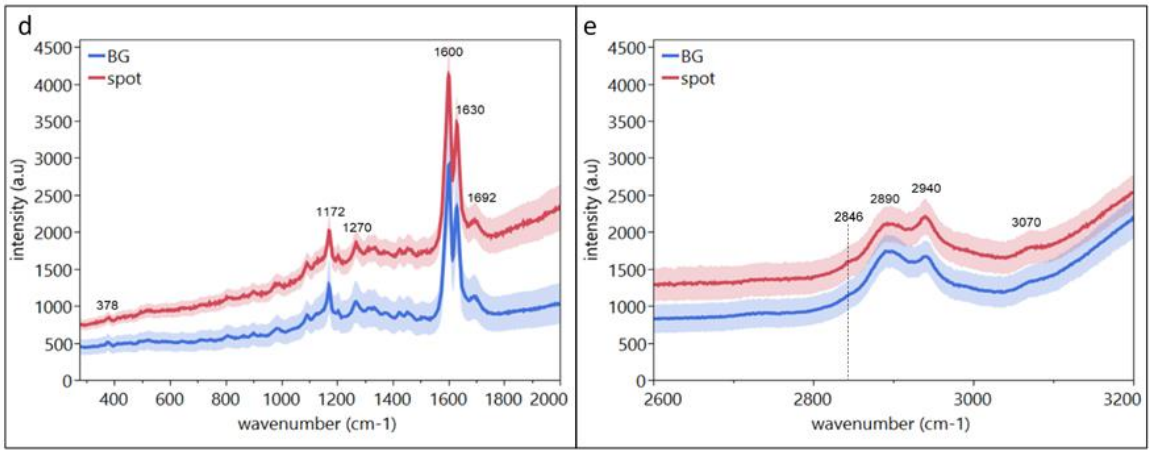
